# Simultaneous task-based BOLD-fMRI and [18-F] FDG functional PET for measurement of neuronal metabolism in the human visual cortex

**DOI:** 10.1101/451468

**Authors:** Sharna D Jamadar, Phillip GD Ward, Shenpeng Li, Francesco Sforazzini, Jakub Baran, Zhaolin Chen, Gary F Egan

**Affiliations:** Monash Biomedical Imaging. Monash University, 770 Blackburn Rd, Melbourne 3800 Australia; Monash Institute for Cognitive and Clinical Neurosciences. Monash University, Wellington Rd, Melbourne 3800 Australia; Australian Research Council Centre of Excellence for Integrative Brain Function. 770 Blackburn Rd, Melbourne 3800 Australia; Department of Electrical and Computer Systems Engineering. Monash University, Wellington Rd, Melbourne 3800 Australia; Department of Biophysics, Faculty of Mathematics and Natural Sciences. University of Rzesow, aleja Tadeusza Rejtana 16C, 35-001 Rzeszów, Poland

**Keywords:** FDG-PET, BOLD-fMRI, glucose metabolism, multimodal imaging

## Abstract

Studies of task-evoked brain activity are the cornerstone of cognitive neuroscience, and unravel the spatial and temporal brain dynamics of cognition in health and disease. Blood oxygenation level dependent functional magnetic resonance imaging (BOLD-fMRI) is one of the most common methods of studying brain function in humans. BOLD-fMRI indirectly infers neuronal activity from regional changes in blood oxygenation and is not a quantitative metric of brain function. Regional variation in glucose metabolism, measured using [18-F] fluorodeoxyglucose positron emission tomography (FDG-PET), provides a more direct and interpretable measure of neuronal activity. However, while the temporal resolution of BOLD-fMRI is in the order of seconds, standard FDG-PET protocols provide a static snapshot of glucose metabolism. Here, we develop a novel experimental design for measurement of task-evoked changes in regional blood oxygenation and glucose metabolism with high temporal resolution. Over a 90-min simultaneous BOLD-fMRI/FDG-PET scan, [18F] FDG was constantly infused to 10 healthy volunteers, who viewed a flickering checkerboard presented in a hierarchical block design. Dynamic task-related changes in blood oxygenation and glucose metabolism were examined with temporal resolution of 2.5sec and 1-min, respectively. Task-related, temporally coherent brain networks of haemodynamic and metabolic connectivity were maximally related in the visual cortex, as expected. Results demonstrate that the hierarchical block design, together with the infusion FDG-PET technique, enabled both modalities to track task-related neural responses with high temporal resolution. The simultaneous MR-PET approach has the potential to provide unique insights into the dynamic haemodynamic and metabolic interactions that underlie cognition in health and disease.

## 1. Introduction

The human brain is a highly metabolic organ requiring a continual supply of glucose to satisfy its energy requirements (1). The neural functions of the brain rely upon a stable and reliable energy supply delivered in the form of glucose through via the blood across the blood-brain barrier and the blood-cerebrospinal fluid barrier (2). The human brain accounts for 20% of the body’s energy consumption at rest (3, 4), of which 70-80% is estimated to be used by neurons during synaptic transmission (5). It is increasingly recognised that access to a reliable energy supply is paramount to maintaining brain health, and early decrements in cerebral glucose metabolism are thought to initiate and contribute to the structural and functional neural changes that underlie age-related cognitive decline (6) and neurodegenerative illnesses (7, 8). In humans, global and regional variations in the cerebral metabolic rate of glucose consumption (CMR_GLC_) can be studied using [18F]-fluorodeoxyglucose positron emission tomography (FDG-PET), which provides a snapshot of glucose utilisation, often averaged over a 30 to 45min period following the injection of the FDG radiotracer.

As cerebral glucose metabolism primarily reflects synaptic transmission, FDG-PET brain imaging has long been used as a proxy for studying human neuronal function in health and disease. However, the FDG-PET bolus administration method has a very limited temporal resolution, representing the integral of the neural (and cognitive) activity occurring during the tracer uptake and PET scanning periods. In addition, PET technology has inherently limited spatial resolution (9), which limits the inferences that can be made regarding the spatial localisation and specificity of metabolic measurements. In contrast, blood oxygenation level dependent functional magnetic resonance imaging (BOLD-fMRI) provides a haemodynamically based surrogate index of neuronal activity with a temporal resolution in the order of seconds (~2sec; or sub-second resolution with advanced multiband MR sequences, e.g., (10) and spatial resolution that can map sub-millimeter neuronal populations, given sufficient field strength and signal-to-noise ratio (11). During the 1990’s the superior temporal and spatial resolution of BOLD-fMRI, together with the fact that it does not require exposure to ionising radiation, led to the BOLD-fMRI method quickly surpassing FDG-PET imaging as the method of choice for studying *in vivo* human brain function.

Despite the apparent ubiquity of the use of BOLD-fMRI for studying human brain function over the last 25-years, the method has a number of shortcomings. The most significant shortcoming of BOLD-fMRI is that the method is not a direct or quantitative measure of neuronal activity. The BOLD-fMRI signal relies upon a complex interplay between metabolic and hemodynamic (blood flow, volume, oxygenation) responses, and the exact relationship between neuronal activity and the measured BOLD signal has not been fully characterised (12). Consequently BOLD-fMRI responses cannot be directly compared across brain regions, subjects and imaging sites, or quantitatively compared in the same individual across time (13). Furthermore, the BOLD signal has been found to contain physiological confounding signals that are non-neuronal, including respiration, heart rate, and arterial responsivity (14). Together these limitations substantially impact the utility of the method for comparing groups of individuals, especially in disease groups (e.g., patients versus healthy controls; aged versus young).

The recent development of simultaneous MR-PET scanning provides the possibility of performing joint spatio-temporal coherence analyses using simultaneously acquired BOLD-fMRI and FDG-PET datasets. Since the potential pitfalls associated with the interpretation of BOLD-fMRI signals are well established, the advantages of combining simultaneously acquired FDG-PET data with BOLD-fMRI may be unclear. The application of joint ICA analytical strategies has the potential to provide enhanced diagnostic and prognostic information compared to a single modality study. The introduction of simultaneous MR-PET allows for the possibility of performing joint analyses which benefit from the signal-to-noise, resolution and sensitivity of BOLD fMRI, together with the interpretability of metabolic measurements and repeatability of semi-quantitative and quantitative FDG-PET.

The technological developments that have led to MR-PET scanners (15) have now made it possible to simultaneously examine changes in blood oxygenation and glucose metabolism at rest, or in response to a stimulus or task, using simultaneously acquired BOLD-fMRI and FDG-PET datasets. BOLD-fMRI acquired simultaneously with PET using a bolus FDG administration has been used to compare sustained (slow) metabolic activity to transient (fast) blood oxygenation changes in rodents (16) and humans (17). Riedl and co-workers (17) compared simultaneous BOLD-fMRI/FDG-PET eyes closed versus eyes open rest in human subjects, and found that local glucose metabolism was highly correlated with BOLD functional connectivity. Wehrl et al. (16) found that simultaneously acquired BOLD and FDG showed different spatial extent and location of peak neural activity using whisker barrel stimulation in the rat. These results suggested that FDGPET measures a more widely distributed neuronal network than BOLD-fMRI, and potentially enables the identification of task-relevant secondary regions that are not detected by BOLD.

Recent advances in FDG infusion protocols (adapted from (18)), together with the improved PET signal detection of dual modality scanners (15) has now made it possible to study dynamic changes in glucose metabolism simultaneously with dynamic changes in blood oxygenation. The method described as ‘functional’ FDG-PET (FDG-**f**PET) involves the radiotracer administered as a constant infusion over the course of the entire PET scan (~90-mins; (19)). In a proof-of-concept study, Villien et al. (19) adapted the constant infusion technique (18) and used simultaneous MR/FDG-**f**PET (but not fMRI) to show dynamic changes in glucose metabolism in response to checkerboard stimulation in the visual cortex with a temporal resolution of approximately one minute. Subsequent studies using simultaneous MR/FDG-**f**PET (20) and BOLD-fMRI/FDG-**f**PET (21) have extended these findings to demonstrate simultaneous BOLD and FDG activity in the visual and motor cortices during alternate visual stimulation (eyes open versus closed) and finger tapping tasks. Interestingly, Hahn et al., (21) found that endogenously-driven stimulation (i.e., eyes open rest and non-paced finger tapping without external behavioural cues) produced task-related CMR_GLC_ and BOLD activity that were independent of each other; in other words, the task-specific changes were not correlated across the two imaging modalities. These results were surprising and in contrast to those reported in the rat (16). In a recent follow-up study, Rischka, Hahn and colleagues (22) showed that endogenously-driven task-related CMR_GLC_ changes can be detected with stimulation periods as low as 2 mins; similar to their previous study, Rischka et al. found no correlation between CMR_GLC_ and BOLD percent signal change.

To date, no study has reported the results of externally-triggered task-based simultaneous BOLD-fMRI/FDG-**f**PET mapping. The investigation of higher-order cognitive functions in the normal human brain, including attention, memory and executive function, involve experimental paradigms and manipulations that result in coupled responses of the concurrent BOLD and FDG signal changes. Conversely, the uncoupling of changes in blood oxygenation and glucose metabolism are thought to underlie many of the cognitive deficits seen in ageing, neurodegenerative diseases and psychiatric conditions. Changes in dynamic regional glucose metabolism may even precede structural, functional and cognitive symptoms in some disease conditions (8). The aim of this study was to test a novel task-based BOLD-fMRI/FDG-**f**PET experimental paradigm that produced concurrent BOLD and FDG signal changes, with a temporal resolution of two seconds and one minute respectively. In this proof-of-concept study, we extended the work of Villien et al. (19), and used a visual checkerboard with varying stimulation durations (Figure 1) and a joint haemodynamic and metabolic analysis approach. We hypothesised that task-based changes in the BOLD signal detected in the brain would be temporally and spatially correlated with task-based changes in FDG metabolism. We identified and investigated joint visual networks and systematically examined the strength and relationship between the concurrently measured BOLD and FDG signals.

**Figure 1:**
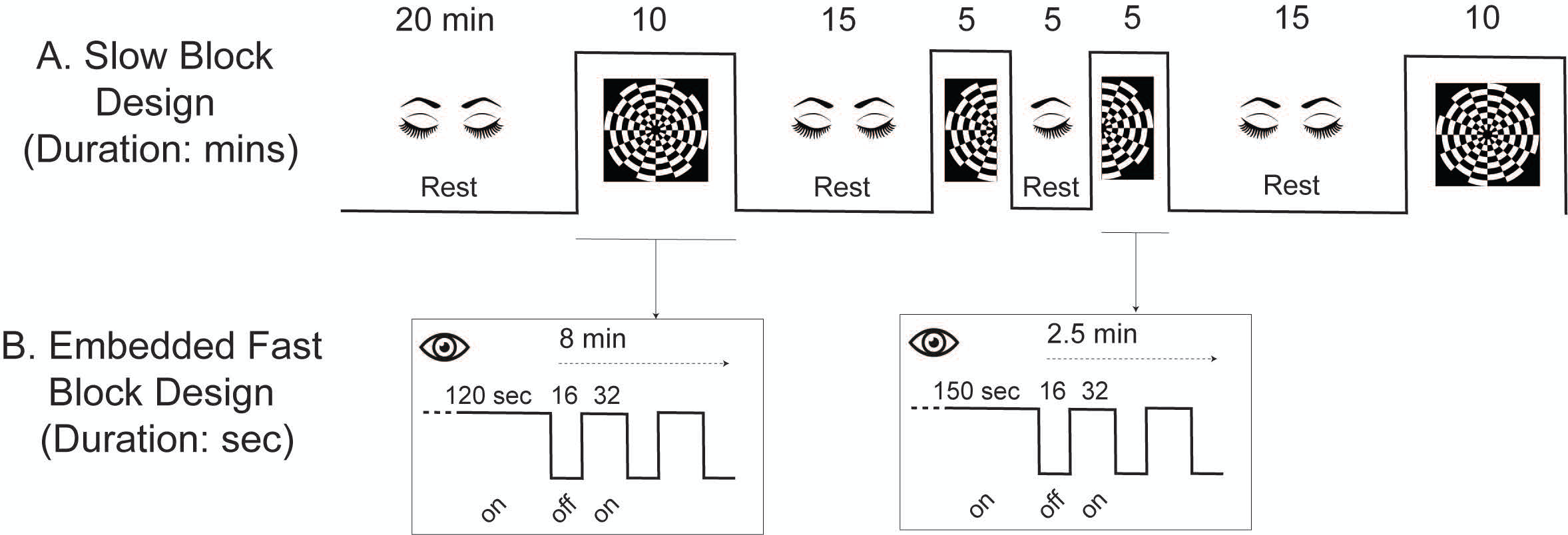
Embedded block design. **A.** Slow block design (durations in minutes) provides contrast for the slow FDG signal (temporal resolution 1-min). Infusion of the tracer started at time 0 in this depiction; to allow the FDG signal to rise to detectable levels, no functional images were acquired during the first 20-mins. During the rest periods, participants rested quietly with their eyes closed. A fast block design (**B**; durations in minutes) was embedded within the checkerboard periods. During the 10-min checkerboard period, the checkerboard was presented for a sustained 120-sec period to allow the FDG signal within the visual cortex to rise above resting levels. During the 5-min half checkerboard periods, the sustained period was 150sec. This was followed by a standard 16sec off/32 sec on alternating design to provide BOLD contrast. During the 16sec off periods, participants viewed a black screen with a fixation cross, and during the 32sec on periods, participants viewed the 8Hz flickering checkerboard on black background.

## 2. Materials & Methods

All methods were reviewed by the Monash University Human Research Ethics Committee, in accordance with the Australian National Statement on Ethical Conduct in Human Research (2007). Data from this study are available on request from the corresponding author.

### 2.1 Participants

Participants (n=10) were aged 18-49 years (mean 29 years), nine female, nine right-handed. Participants had 13-26 years of education (mean 17.6 years), and had normal or corrected-to-normal vision. Participants were screened for diabetes, personal or family history of a neurological neurodegenerative condition, claustrophobia, and non-MR compatible implants; women were screened for current or suspected pregnancy. Prior to the scan, participants were directed to consume a high protein/low sugar diet for 24 hours, fast for 6 hours and drink 2-6 glasses of water. Blood sugar level (BSL) was measured using an Accu-Chek Performa (model NC, Mannheim, Germany); all participants had BSL below 10mML (max 5.8mML).

### 2.2 Stimuli and Tasks

We developed an embedded block design to produce signal contrast for both fast BOLD-fMRI and slow FDG-**f**PET measurements (Figure 1). The slow on/off design (Figure 1A) was adapted from Villien et al. (19) and provided FDG-**f**PET contrast. The fast on/off design (Figure 1B) was embedded within the stimulation periods and provided BOLD-fMRI contrast.

Participants rested with eyes closed during the initial 20 minute block of non-functional scans (localiser, T1, etc). At 20 mins, a 10 min stimulation period was presented in an embedded 32/16-sec on/off design. The first 120 seconds of the block was a sustained ‘on’ period to allow the FDG-**f**PET signal to rise from resting levels. Following that 120 sec period, the 32/16 sec on/off design was implemented. During the ‘on’ periods, the visual stimulus was a circular checkerboard of size 39cm (visual angle 9°) presented on a black background. The checkerboard flickered (i.e., fields alternated black/white) at 8Hz. During the ‘off’ periods, participants rested eyes open while viewing a white fixation cross of size 3cm (visual angle 0° 45’) presented on a black background. A 15 min eyes closed rest period followed, then 5 mins of left hemifield stimulation (150 sec on, followed by 32/16 sec on/off), 5 mins of eyes closed rest, and 5 mins of right hemifield stimulation. Note that participants were instructed to open both eyes during hemifield stimulation. Following 20 mins of eyes closed rest, a final 10 min full checkerboard stimulation was presented.

### 2.3 Procedure

Participants were cannulated in the vein in each forearm with a minimum size 22-gauge cannula, and a 10mL baseline blood sample was taken at time of cannulation. For all participants, the left cannula was used for FDG infusion, and the right cannula was used for blood sampling. Primed extension tubing was connected to the right cannula (for blood sampling) via a three-way tap.

Participants underwent a 90-minute simultaneous MR-PET scan in a Biograph 3Tesla molecular MR (mMR) scanner (Siemens, Erlangen). Participants were positioned supine in the scanner bore with head in a 16 channel radiofrequency (RF) head coil. Visual stimuli were viewed through a mirror placed on the RF coil which projected to a 32-inch BOLDscreen (Cambridge Research Systems, UK) MR-compatible LCD screen. Consistent with the protocol reported by Villien et al. (19), [18F]-Fluorodeoxyglucose (average dose 97.0 MBq) was infused over the course of the scan at a rate of 36mL/hr using a BodyGuard 323 MR-compatible infusion pump (Caesarea Medical Electronics, Caesarea, Israel). One participant received a dose of 39 MBq due to technical error. Infusion onset was locked to the onset of the PET scan (see MR-PET Scan Protocol, below).

Plasma radioactivity levels were measured throughout the duration of the scan. At 10 mins post-infusion onset, a 10mL blood sample was taken from the right forearm using a vacutainer; the time of the 5mL mark was noted for subsequent decay correction. Subsequent blood samples were taken at 10 min intervals for a total of 10 samples for the duration of the scan. The cannula line was flushed with 10mL of saline after every sample to minimise line clotting. The mean plasma radioactivity concentration over time is presented in the Supplementary Results.

### 2.4 MR-PET Scan Protocol

PET data was acquired in list mode. Infusion of FDG and PET data acquisition started with Ultrashort TE (UTE) MRI for PET attenuation correction. To allow the PET signal to rise to detectable levels, non-functional MRI scans were acquired in the first 20-mins following infusion onset. These scans included T1 3D MPRAGE (TA = 7.01mins, TR=1640ms, TE=2.34ms, flip angle = 8°, FOV=256×256mm^2^, voxel size = 1×1×1mm^3^, 176 slices; sagittal acquisition), T2 FLAIR (TA=5.52mins; data not reported here), gradient field map (TA=1.02mins, TR=466ms, TE_1_=4.92ms, TE_2_=7.38ms, voxel size=3×3×3mm^3^, FOV=190mm, 44 slices), and ASL (TA=3.52min, data not reported here). For the remainder of the scan, seven consecutive blocks of T2*-weighted echo planar images (EPIs) were acquired (TR=2450ms, TE=30ms, FOV=190mm, 3×3×3mm^3^ voxels, 44 slices, ascending interleaved axial acquisition). The duration of each T2* block was determined by the stimulus block duration (Figure 1A): block 1 full checkerboard (TA=10.02min, 242 volumes), block 2 eyes closed rest (TA=15.01min, 364 volumes), block 3 half checkerboard (TA=5.05min, 121 volumes), block 4 eyes closed rest (TA=5.05min, 121 volumes), block 5 half checkerboard (TA=5.05min, 121 volumes), block 6 eyes closed rest (TA=20.02, 487 volumes), block 7 full checkerboard (TA=10.02min, 242 volumes).

#### PET Image Reconstruction

We examined both static FDG-PET and dynamic FDG-fPET results. For the static FDG-PET analysis, PET list mode data were reconstructed in one 95-minute image. For the dynamic FDG-**f**PET analysis, PET list mode data were binned into 95 one minute frames. All data were reconstructed offline using the e7tools software (provided by Siemens) and corrected for attenuation using pseudoCT (23) and motion (24). Note that this motion correction algorithm corrects across the entire PET experiment, unlike other methods that only correct during T2* MR acquisition. Ordinary Poisson - Ordered Subset Expectation Maximization (OPOSEM) algorithm with Point Spread Function (PSF) modelling (25) was used with 3 iterations, 21 subsets and 344×344×127 (voxels size: 2.09×2.09×2.03 mm^3^) reconstruction matrix size. A 5-mm 3-D Gaussian post-filtering was applied to the final reconstructed images.

### 2.5 Data Analysis

We conducted a data-driven analysis using an independent component analysis (ICA) strategy, because we chose to not impose strong a priori hypotheses on the form and shape of the expected FDG-fPET task-based activation response. We did not want to constrain the search for task-related activity on the basis of a fixed model-based hypothesis, as is required in a General Linear Model (GLM) analysis. Furthermore, as we were interested in developing and testing the novel experimental paradigm (hierarchical block design) to simultaneously measure high temporal resolution BOLD-fMRI and FDG-fPET, we have conducted a semi-quantitative rather than quantitative analyses of the FDG-PET images.

Image analysis was conducted in four steps. An independent component analysis was conducted on (i) BOLD-fMRI, (ii) static FDG-PET, and (iii) dynamic FDG-**f**PET data separately. Finally (iv) the ICA components for visual cortex for BOLD-fMRI and dynamic FDG-**f**PET (from (i) and (iii)) were entered into a joint ICA to estimate components that share the same mixing parameters. For each step, the components showing primary visual cortex activity are reported below. The results for the remaining components are reported in the Supplementary Results.

#### 2.5.1 MRI Data

The four task blocks (two full and two half checkerboard blocks) for all subjects underwent standard fMRI preprocessing in FSL, including B0 unwarping, motion correction, and high pass filtering (0.01Hz). Filtered images were entered into a first-level independent component analysis with automatic estimation of the number of components using MELODIC (26). Temporal trends from manually classified noisy components were regressed out from the data. In order to normalize the cleaned data to the MNI space, the following steps were performed. Firstly, the T1-weighted image was non-linearly registered to the MNI template using the symmetric normalization algorithm in ANTs. Then, the first volume of the cleaned file was linearly registered to the T1w image using 6 degrees of freedom. The last step was to apply all the transformations from the previous two points to the 4D cleaned file. Finally, the normalized cleaned file was smoothed with a 5mm FWHM Gaussian kernel.

Smoothed data for all tasks and for all subjects were used to compute a multi-session, multi-subject ICA using GIFT v4.0b (27), implemented in MatLab (Natick, Massachusetts USA), with the number of independent components set to seven. This number of components was chosen as it resulted in a stable decomposition without splitting of known brain networks across multiple components. The independent component corresponding to the visual network was retained. A voxel-wise t-test was calculated for this component using the four back-reconstructed Z-maps (one for each task block separately) to obtain one mean visual activation map per subject. The four Z-maps were retained for the joint analysis.

#### 2.5.2 PET Data

Note that in this analysis we examined change in FDG uptake across the experiment, we did not calculate CMR_GLC_. Visual inspection of the rigid body registration of the FDG-PET images to MNI space highlighted the poor quality of the subject-level FDG-PET image registrations to the T1 weighted MR images. An FDGPET template was therefore constructed that enabled accurate alignment of each subject’s FDG-PET image to the template and to their T1 weighted MR image. The study-specific PET template was created using six randomly chosen subjects from the cohort. Firstly, five subjects were linearly registered to one reference subject using 12 degrees of freedom registration. The mean (static) image for each subject was input into ANTs (antsMultivariateTemplateConstruction2 http://stnava.github.io/ANTs/) to generate the template. This function uses the symmetric group-wise normalization (SyGN) algorithm to provide an optimal and un-biased population template (28, 29).

For the static FDG-PET analysis, the static motion-corrected FDG-PET images for all subjects were registered to the study specific PET template and merged together into a 4D image. Normalisation was performed using the cerebellum as a reference region (30). Seven independent components were extracted using GIFT, and the primary visual network component was manually identified and aligned to MNI space for visualisation.

For the dynamic FDG-**f**PET analysis, grey matter masks were generated for each subject using FreeSurfer and the T1 and T2 structural images. The first ten frames (10 mins) of each subject were removed because of the low number of coincidence counts at the beginning of the infusion procedure. The remaining 4D volumes and grey matter masks were registered to the PET template. The total spatial smoothing applied to each volume was equivalent to 6mm FWHM including the Gaussian smoothing applied during reconstruction. The 4D volume of each subject was normalized as follows in order to remove the inter-subject variance in the non-task related changes in the FDG-PET signal, as follows: (1) a 3^rd^ order polynomial regression was used to fit the time activity curve of the average whole brain grey matter uptake for each subject; and (2) the time series of each voxel was rescaled by division of the grey matter trend computed in step (1), resulting in region specific relative FDG signal uptake and neural activity curves.

The normalized 4D volume represented the dynamic relative glucose uptake (time-activity) map. The volumes for all subjects were temporally concatenated to form an augmented spatiotemporal matrix for group independent component analysis (ICA). The FastICA toolbox (31) was used in Matlab to undertake the spatial ICA analysis. The number of independent components was set to seven to match the MRI analysis method, and the data dimension was reduced to seven by using principal component analysis (PCA) methods with 76.9% of the variance reserved. The component corresponding to the visual network was manually identified and dual regression (32, 33) was used to generate subject-specific Z-score maps. These maps were retained for the joint analysis.

#### 2.5.3 Joint Analysis

Joint ICA (34) was performed using the Fusion ICA Toolbox (FIT, v20d) implemented in MatLab. The single-subject Z-maps from the BOLD-fMRI and the dynamic FDG-**f**PET analyses were concatenated into two 4D files and entered into a joint ICA with the number of extracted components set to three. The joint ICA algorithm proceeded as follows: (1) the spatial component maps for both the fMRI and fPET analyses were normalised to have the same average variance (sum-of-squares) when computed across all subjects and voxels; (2) the spatial maps for each modality and for each subject were vectorised and merged into one NxM matrix, with N equal to the number of participants and M to the number of voxels multiplied by two; (3) PCA was subsequently used to reduce the dimensionality of the data (to three dimensions) and retain the key features from both modalities; and (4) the extended-infomax algorithm (35) was used to decompose the reduced feature matrix to three maximally independent components, for each modality. Here we report the results for the first joint component, and see the Supplementary Results for the other two components.

#### 2.5.4 Probabilistic Anatomy Mapping

In order to examine the spatial characteristics of the BOLD-fMRI and the static and dynamic FDG-PET results, we used statistical probabilistic anatomical mapping (SPAM) (36) implemented in the SPM12 Anatomy Toolbox (v2.2c). The probabilistic cytoarchitectonic maps are based on an observer-independent cytoarchitectonic analysis in a sample of ten human post-mortem brains, and provide stereotaxic information on the location and variability of cortical areas in the Montreal Neurological Institute (MNI) reference space (37). The visual network component maps from the ICA analyses of the BOLD-fMRI, static FDG-PET, and dynamic FDG-fPET datasets were entered into the SPAM analysis, thresholded to Z>2, k>200 voxels, and the anatomical mapping probability threshold set to p<0.05.

#### 2.5.5 Similarity analysis

To quantify the similarity between the visual network maps (one for fMRI and one for fPET) extracted from the joint ICA analysis, we calculated the Dice Similarity Score (DSS) coefficient. The DDS was computed by applying fifty Z-score threshold values, from the 95^th^ to the 100^th^ percentile with step size of 0.1, to binarize (0 or 1 in all the voxels which were below or above the threshold, respectively) for the visual network component maps from both modalities (38). To ensure the DSS did not harshly penalise near misses, particularly due to image alignment and smoothing pre-processing, a morphological dilation of 1 voxel was applied to the binary maps when calculating the number of true positives (TP) in the DSS equation (see equation 1, dilated mask denoted by δ).

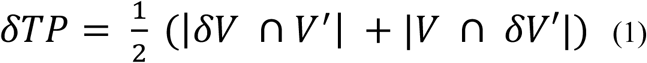

*V* and *V*′ were the fMRI and fPET binary visual maps respectively. The DSS between each pair of binarized maps was computed as shown in equation 2.

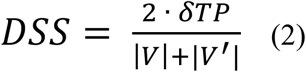

## 3. Results

The image data for both modalities were analysed separately, and then jointly, using a joint ICA to calculate the components of maximal correlation. Here, we report the results for the primary visual network extracted from each analysis. This network was the first component obtained from each analysis, in a manner consistent with the experimental design (Figure 1). The results for the other components estimated in each analysis are reported in the Supplementary Results.

The static FDG-PET analysis revealed activity predominantly in bilateral occipito-parietal areas (Figure 2B; Supplementary Table 1), including primary visual cortex (V1), secondary visual cortex (V2), visual areas V3 and V4, fusiform gyrus, intraparietal sulcus, superior and inferior parietal lobules and cerebellum. The dynamic FDG-fPET analysis revealed a similar pattern of results (Figure 2C, Supplementary Table 1), with activity primarily in the visual cortex, including V1, V2, V3, fusiform gyrus, and superior and inferior parietal lobules. Lastly, the conventional BOLD-fMRI analysis (Figure 2A, Supplementary Table 1) showed a similar pattern of results, with activity seen in the same visual areas (V1, V2, V3, V4, fusiform gyrus), cerebellum, and superior parietal lobule. The results of the dynamic FDG-**f**PET and BOLD-fMRI analyses confirmed that the embedded block design yielded robust activity within the expected regions of visual cortex. The static FDG-PET and dynamic FDG-**f**PET results were also similar, except for the addition of activity that was identified in the cerebellum and in the intraparietal sulcus in the static, but not in the dynamic analysis. This discrepancy may be attributed to the additional statistical power in the static analysis due to signal averaging that enabled the identification of smaller effects.

**Figure 2:**
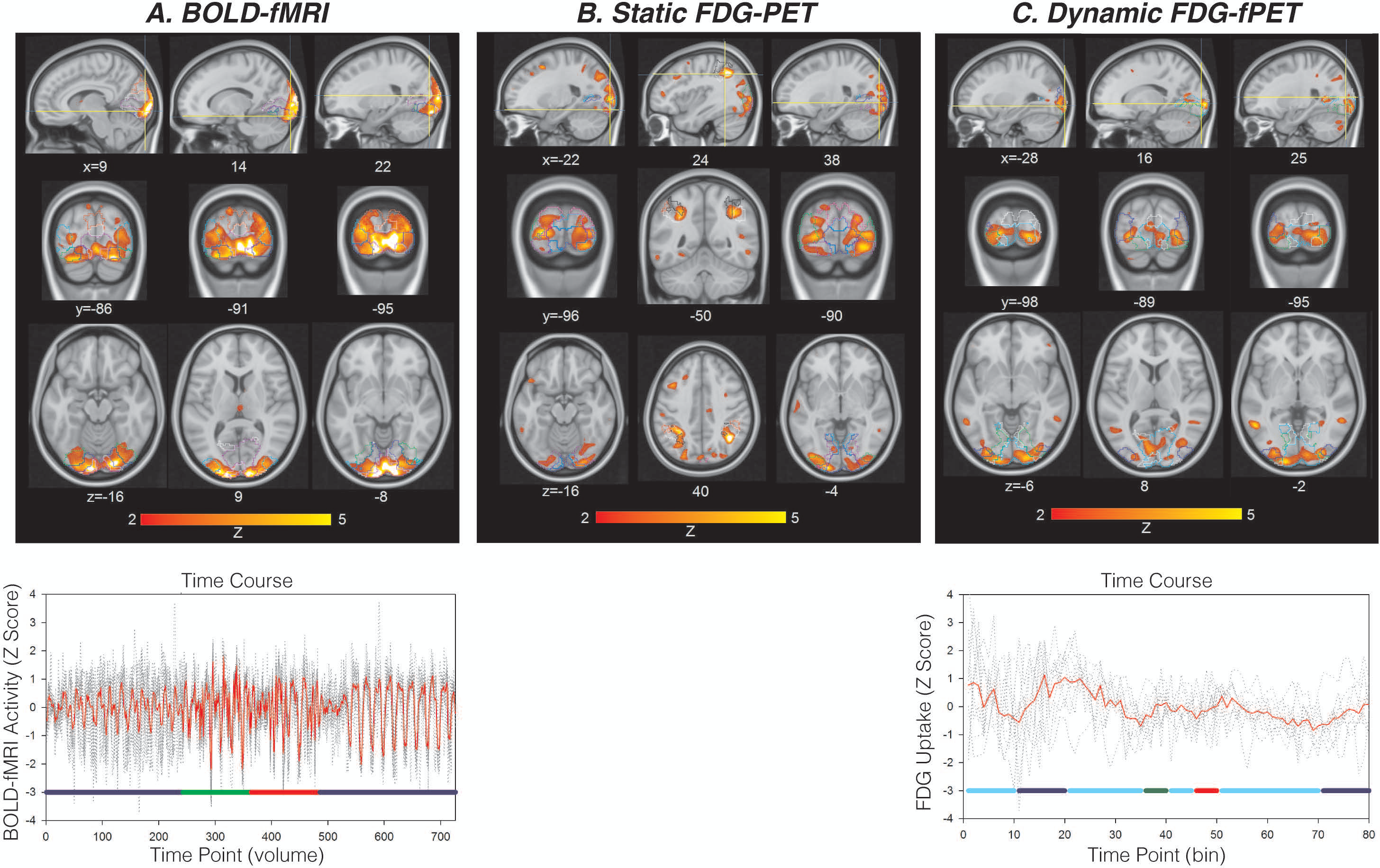
**A.** (Top) BOLD-fMRI independent component 1 showing the visual network. (Bottom) Timecourse for BOLD-fMRI independent component 1. **B**. Static FDG-PET independent component 1 showing the visual network. **C.** (Top) Dynamic FDG-**f**PET independent component 1 showing the visual network. (Bottom) Timecourse for dynamic FDG-**f**PET independent component 1. Coordinates are MNI. Statistical Parameter Anatomical Maps are shown in coloured outline on transverse and coronal images. For timecourses, red line shows mean across subjects, individual subject timecourse shown in dotted lines. Coloured bars indicate the timing of the experimental design; light blue: rest; dark blue: full checkerboard; green/red: half checkerboard. See Supplementary Table 1 for coordinates and statistical results.

The joint ICA of BOLD-fMRI and dynamic FDG-fPET data identified one joint visual network component (Figure 3A&B, Supplementary Table 2). The BOLD-fMRI component showed a larger extent of activity than the FDG-fPET component, but both showed activity in qualitatively similar regions across the visual cortex. To quantify the extent of similarity between the joint components, we calculated a similarity matrix using Dice similarity coefficients (Figure 3C). This analysis, conducted across the whole brain for the first joint fMRI and fPET component (visual network, Figure 3A&B) confirmed that the peak of activity in each map is highly spatially concordant for the two modalities.

**Figure 3:**
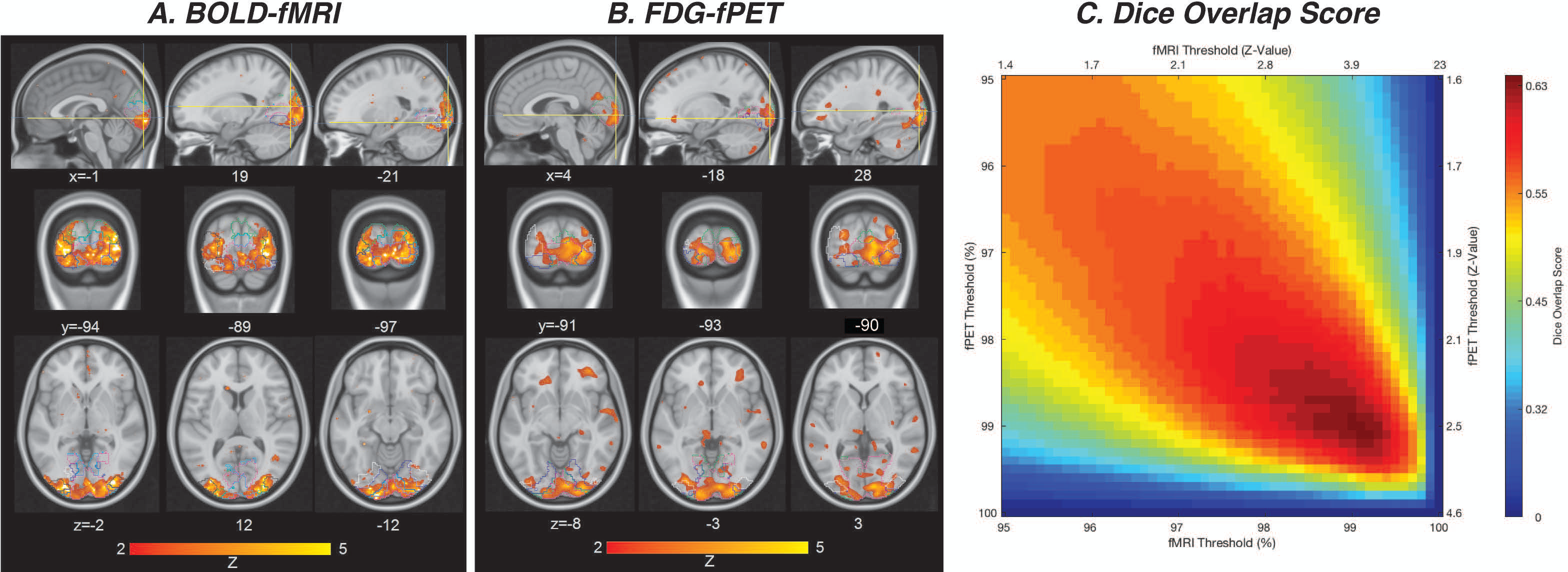
**A & B**Joint ICA results; first component pair showing activity in the visual network. Coordinates are MNI. Statistical Parameter Anatomical Maps are shown in coloured outline on transverse and coronal images. See Supplementary Table 2 for coordinates and statistical results. **C.** Dice overlap score for statistical similarity between BOLD and FDG components identified in the joint ICA, calculated across the whole brain. Colourbar indicates Dice coefficient strength.

## 4. Discussion

In this brain imaging study, we investigated the relationship between simultaneously acquired measures of human brain activity. Extending the work of Villien et al. (16), we developed and tested a novel experimental paradigm to induce neural activity in the visual cortices with high temporal resolution using simultaneously acquired BOLD-fMRI and dynamic FDG-**f**PET. In the **f**PET analysis we focused on measurement of relative change in FDG uptake (glucose metabolism) but we did not quantify CMR_GLC_ (similar to previous research, e.g., Riedl et al.(17)). Our results demonstrate that the hierarchical block design with fast and slow visual stimuli enabled both modalities to track task-related neural responses with improved temporal resolution than obtained in bolus FDG-PET studies. The most prominent brain network identified in both modalities was located in the primary visual cortex network which is known to be associated with visual stimulation. Importantly, these results show that it is possible to examine transient metabolic responses to task stimuli using a hierarchical block design and a slow infusion dosing protocol (18, 19) with higher temporal resolution than has previously been possible using bolus protocols and simple alternating on/off experimental designs.

Our primary motivation for using an independent components analysis strategy was to conduct a data-driven analysis. Traditional approaches like the GLM provide a constrained model-based analysis, which makes assumptions about the trial-by-trial shape and variation of the neural response. Analysis of BOLD-fMRI data relies upon the balloon model of the relationship between the neuronal activity and haemodynamic response, as well as the canonical haemodynamic response function. Analogous models linking neural activity to FDG uptake and trial-by-trial metabolic response functions are not yet available in human MR-PET studies.

The small number of published studies that have examined the relationship between resting-state BOLD-fMRI and FDG-PET have reported that there is little overlap between the networks obtained in BOLD and FDG, with the FDG components showing greater left-right connectivity and the BOLD networks showing greater anterior-posterior connectivity (e.g., (39, 40)). We decided to undertake an independent components analysis strategy for these reasons, and because the data-driven approach has the potential to uncover a greater diversity of task-related activity than the constrained GLM approach, it was more suited to address the exploratory goals of the current study.

It is important to understand the assumptions that underlie the observed relationship between the BOLD and FDG data indicated by the joint analysis. Spatial independent components analysis (ICA) assumes that the signal represents a set of spatially independent networks (components), each with associated time-courses (27, 41). The analytical approach constrains each voxel within a component to have the same temporal fluctuations of blood oxygenation in BOLD-fMRI, or of glucose uptake in FDG-**f**PET. Thus, each independent component calculated in the group analysis represents a temporally coherent brain network of either haemodynamic or metabolic ‘connectivity’. The joint-ICA approach is a second-level feature-based analysis, which concatenates data from the two modalities to estimate a joint unmixing matrix *across subjects* (42). The algorithm assumes that the sources of neural activity associated with the two data sets modulate the same way across subjects (43), but it does not make assumptions regarding the linearity (or non-linearity) of the relationship between the underlying physiological processes under investigation. Thus, we can conclude that within the visual network components identified in the joint analysis, the strength of the positive relationship between the FDG and BOLD measures varies systematically across individuals. The FDG infusion protocol we used in this experiment has produced significantly improved temporal resolution compared to previous studies (39, 40, 44). The improved FDG temporal resolution, together with the data fusion analytical approach, provides a powerful framework for exploring the relationship between task-evoked local changes in blood oxygenation and glucose metabolism. Until now this has proved elusive using univariate correlational approaches (21).

Our results complement and extend the small number of existing simultaneous BOLD-fMRI/FDG-PET studies reported to date. Consistent with Riedl et al. (17) and Wehrl et al. (16), we found that blood oxygenation changes in the tissue were related to changes in glucose metabolism in the visual network (however Hahn et al., 2018 (21) found no relationship between glucose metabolism and functional connectivity in the visual and motor cortices using simultaneous BOLD-fMRI/FDG-**f**PET). Riedl et al. (17) showed that local glucose metabolism was highly correlated with BOLD functional connectivity using a simultaneous BOLD-fMRI/FDG-PET with bolus design comparing eyes open versus closed rest, consistent with results obtained in non-simultaneously acquired data (39, 44-46). In a follow-up analysis of Riedl et al.’s data, Savio et al. (40) showed a modest statistical spatial similarity between the modalities: in comparison to BOLD resting-state networks, networks extracted from the FDG data were less differentiated and showed smaller clusters of activity. In contrast, in a simultaneous BOLD-fMRI/FDG-PET with bolus injection in the rat, Wehrl et al. (16) found that box-whisker stimulation activated a qualitatively more distributed network in FDG than BOLD. In the current study, comparison of the joint components showed that at high statistical thresholds, the clusters around the peaks of activity were statistically similar in spatial location. In other words, across the whole brain, clusters in the joint visual FDG-BOLD component showed good (but not excellent, maximum Dice coefficient ~0.65) statistical similarity when the FDG and BOLD thresholds were matched.

The degree of spatial mismatch observed between the FDG and BOLD signals is likely to be attributable to the different physiological bases of each brain imaging methodology. The FDG signal primarily arises in the neural tissue and is more directly related to neuronal activity than the BOLD signal, which can be affected by post-capillary down-stream effects in the cerebral vasculature. In particular, the characteristics of the signal-to-noise ratio differs between the two methods, which not only affects the statistical power of the methods to detect effects, but also the proportion of voxels/clusters that exceed statistical threshold. The on/off design did differ between the fast (fMRI) and the slow (fPET) components. Subjects were instructed to have their eyes closed during the extended (10min) rest periods, and their eyes open and fixated during the embedded (16sec) rest periods. However, we expect that this difference in the fMRI and fPET rest conditions is unlikely to account for the spatial mismatch between the activation maps for the two modalities. In their comparison of eyes open and eyes closed rest, Riedl et al., used a conjunction analysis to show that the FDG and BOLD maps that compared eyes open versus eyes closed showed strong overlap in the visual regions. Furthermore, they computed a spatial similarity coefficient that showed no significant difference between the eyes closed and open contrast for the two modalities. Their results are compatible with our contention that the difference in the rest condition in our experimental design does not account for the spatial mismatch between the FDG and BOLD visual activity maps.

The highest temporal resolution for detection of FDG changes in the brain is of primary interest and an important area for future research. The experimental task design for the fMRI acquisition included an 18 sec rest period during the visual paradigm. However, we are uncertain whether the glucose metabolism returned to baseline during the rest period, as we do not have the necessary temporal resolution nor the sensitivity when measuring the FDG-PET signal to resolve this question. However, the FDG-PET signals in the visual cortex would be expected to be at least two thirds higher during the blocks of visual stimulation than during the between-block rest periods.

Finally, the spatial resolution of PET is inherently limited, which affects the spatial specificity of the results^*^. Our results contribute significantly to the literature concerning simultaneous and non-simultaneous MR-PET that suggests a relationship between glucose metabolism measured using FDG-PET and the haemodynamic response in BOLD-fMRI (16, 17, 19, 39, 44-46, 48).

Simultaneous BOLD-fMRI/FDG-PET and **f**PET methodology is still in its infancy, with most existing studies examining intrinsic connectivity with bolus (resting-state BOLD-fMRI/FDG-PET: (17, 40, 48). Our study is most directly comparable to Hahn et al.’s (21) and Rischka et al.’s (22) study, the only other published studies that used task-based simultaneous dynamic BOLD-fMRI/FDG-**f**PET. Hahn et al. studied self-paced eyes-open versus eyes-closed and finger tapping, and found that regions showing BOLD-fMRI response to each task did not show a concomitant change in CMR_GLC_. Rischka et al. studied video viewing and eyes-open finger tapping and also found that there was no significant correlation in the percent signal change between fPET and fMRI. Hahn and colleagues’ finding that there is no relationship between fPET and fMRI are in contrast to the current results, as well as evidence from resting-state BOLD-fMRI/FDG-PET studies (e.g., (16, 17, 40)). Our novel experimental design may have been a crucial factor that enabled us to detect hitherto unreported covariances between BOLD and FDG. This was achieved by time-locking the stimulus-response relationship in the two modalities in a tightly-controlled embedded (hierarchical) block design. However, this would suggest that endogenously-triggered task-based neuronal activity shows a neurovascular-metabolic relationship that is different to resting or exogenously-triggered task-based neuronal activity, which seems rather unlikely. It is more likely that the data fusion analysis provided more power to detect latent relationships between the FDG and BOLD data than Hahn and colleagues’ univariate correlational approach.

The conflicting interpretations suggested by existing BOLD-fMRI/FDG-PET studies may be addressed by future simultaneous MR-PET studies that examine the coupling of cerebral blood flow and metabolism. There is a strong relationship between capillary blood volume and glucose uptake (49), and since capillary diameter (and thus blood volume) is a modulator of cerebral blood flow (50), coupling between FDG and BOLD measured responses may be expected. However, FDG-PET and BOLD-fMRI do not exclusively measure cerebral rate of glucose metabolism and cerebral blood flow. The FDG image comprises signals from blood and the tissue compartment in each voxel. The BOLD signal captures downstream venous blood oxygenation changes as well as blood flow fluctuations that may be due to causes in addition to neural activity. Accordingly, the spatial and/or temporal correlations between the two signals are not assured to be always related. Simultaneous MR-PET, together with advanced data fusion analytical techniques, has the potential to investigate and elucidate the assumptions associated with coupling between cerebral metabolism and cerebral blood flow in the healthy brain as well as in disease. Further studies, involving quantitative or calibrated fMRI techniques together with MR-PET (51), have the potential to offer additional insights into the shift between neural energy sources and the relative utility of glucose and oxygen cerebral metabolic processes.

Within multimodal imaging neuroscience, there are four main methods for data fusion (52). Many studies use a qualitative visual comparison (e.g. (39)) which can be very informative but do not allow quantitative inferences to be made. Most existing BOLD-fMRI/FDG-PET studies use a data integration approach, where data from each modality is analysed separately and compared at the second-level using correlation (e.g., (16, 21, 45, 46, 48)) or similarity metrics (e.g., (40)). A data integration approach is useful for examining broad relationships between data types, but does not fully capitalise on the shared information between the modalities. Asymmetric data fusion is common in EEG-fMRI studies (53), and uses information from one modality to constrain the other (e.g., PET-constrained MR regions-of-interest; (17)). This approach can impose unrealistic assumptions (i.e., FDG and BOLD measure fundamentally different processes, and a response in one modality may not show an analogous response in the other) and also can bias the analyses that compare the two datasets. A data fusion approach, as used here, treats each modality equally in order to take advantage of the joint information in each data set. As reviewed by Calhoun & Sui (52), data fusion approaches have shown improved ability to uncover latent relationships between data types, which in turn have been useful for radiographic biomarker (FDA-NIH Biomarker Working Group) (54) discovery studies in psychiatry. Thus, it is likely that our powerful data fusion approach to BOLD-fMRI/FDG-fPET has allowed us to identify relationships between task-based BOLD and FDG data that previously have not been captured by a univariate correlational analysis, as used by Hahn et al. (21).

Our study is one of the few existing studies using simultaneous MR-PET to study relationships between BOLD and glucose metabolism (16, 17, 21, 40, 48) and only the third study to examine task-based responses (22). Across imaging neuroscience, simultaneous acquisition of multimodal data is preferable to consecutive acquisitions, as they minimise inter-session variability in alertness, fatigue, behaviour, practice effects and task engagement. Simultaneous acquisitions are also preferred as data is optimally registered, both spatially and temporally (i.e., BOLD responses are intrinsically time-locked to FDG responses). However, substantial advancement in methodology is usually required in multimodal acquisitions, including removal of artefacts in one modality that are introduced by the other, as in the case of simultaneous EEG-fMRI. In the case of simultaneous MR-PET, the mutual interference of MR and PET technologies has represented a major technical challenge in the development of multimodal scanners, image processing and analysis techniques (reviewed in Chen et al., in press). Improvements in PET image reconstruction, attenuation and motion correction are the focus of ongoing development.

While our data fusion procedure was a powerful method for uncovering latent relationships between the BOLD and FDG data (52), like previous studies our analysis did not fully take advantage of the simultaneous nature of the acquisition. Rather we have performed direct data fusion on high-dimensional summary measures. Our analysis approach (and those of existing BOLD-fMRI/FDG-PET studies) require a number of parameter selections, including smoothing factors and pre-specifying the number of components to be estimated by ICA. In our approach, we selected the number of components (seven) based on previous BOLD-fMRI studies and applied this to data from both modalities. While this number of components yielded a stable decomposition in the BOLD data, the results from the current and previous studies indicate that FDG-PET data may require a lower number of components to be estimated. In this study, there was some evidence of separation (or ‘splitting’) of FDG-fPET components (see Supplementary Figures 3 and 4), consistent with previous resting-state BOLD-fMRI/FDG-PET analyses (39, 40). Functional connectivity estimated from resting-state BOLD-fMRI has benefited from the evidence gathered in the thousands of published studies since its introduction in 1995 (55), and thus a great deal of further work is required to optimise the analysis protocols of metabolic connectivity using fPET.

## 5. Conclusions

In this study, we have presented a novel experimental design for the measurement of task-evoked changes in brain activity with high temporal resolution that uses simultaneous BOLD-fMRI/FDG-**f**PET imaging to capture metabolic and haemodynamics responses. We showed that this approach yields task-related activity in both modalities during simultaneous BOLD-fMRI/FDG-**f**PET acquisition. While studies of the resting-state of the brain are important for understanding quiescent neural activity, studies of task induced brain activity remain the cornerstone of cognitive neuroscience for unravelling the spatial and temporal brain dynamics of cognition in health and disease. The simultaneous MR-PET approach has the potential to provide unique insights into the dynamic haemodynamic and metabolic interactions that underlie impaired cognition in psychiatric and neurodegenerative illnesses.

## 6. Acknowledgements

We thank Richard McIntyre, Alexandra Carey and Jason Bradley for assistance with the implementation of the slow-infusion technique and scanning protocol. We thank Edwina Orchard, Irene Graafsma and Disha Sasan, and the staff at Monash Biomedical Imaging for assistance with data collection.

Jamadar is supported by an Australian Council for Research (ARC) Discovery Early Career Researcher Award (DECRA DE150100406). Jamadar, Ward and Egan are supported by the ARC Centre of Excellence for Integrative Brain Function (CE114100007). Chen and Li are supported by funding from the Reignwood Cultural Foundation.

## 8. Author Contributions

SJ, ZC and GE designed the study; all authors contributed to analysis design; FS, SL, JB analysed the data; all authors contributed to manuscript preparation.

## Footnotes

* In some instances, even when BOLD and FDG images are simultaneously acquired, the images may not be tightly registered at acquisition, as the centre of the PET FOV may not align with the isocentre of the MR scanner 47. Catana C, *et al.* (2011) MRI-assisted PET motion correction for neurologic studies in an integrated MR-PET scanner. *Journal of Nuclear Medicine* 52(1):154-161.; furthermore, the PET FOV is larger than MR FOV. Such hardware characteristics may require the application of a transformation matrix that may impact the registration of the final images. This was not the case with data acquired on our scanner, and registration checks during pre-processing confirmed that T1 MRI and PET UTE scans were well registered at the time of acquisition.

## Supplementary Information

### Supplementary Results

#### Plasma Radioactivity Results

Supplementary Figure 1 shows the interpolated plasma radioactivity concentration over time for subjects included within the experiment. The total counts value detected by the counting tube were decay corrected to obtain the activity of the plasma at the blood collection time.

Due to the issues with blood collection, plasma samples were not obtained for one patient and only three measurements were performed for two another. Two volunteers received significantly lower overall dose (39MBq and 73.3 MBq, respectively) than the remaining participants due to technical issues with the pump which stopped infusion of the radiopharmaceutical during the scan.

The averaged radioactivity concentration constantly increases over the time with the lowest relative slope at the end of the acquisition (Supplementary Figure 1). It shows that equilibrium in the plasma was not reached during the experiment. However, the analysis of the individual concentration curves reveals that for subjects for whom the pump failed the equilibrium was reached for a approx. 30 minutes and decreased after that. A limitation of our study is that plasma radioactivity did not reach equilibrium during the PET acquisition, consistent with previous studies using the constant infusion approach (1, 2). We believe, as it is shown in (3, 4), a bolus plus continuous infusion strategy will allow plasma radioactivity to reach the equilibrium much faster than the approach used in this paper.

**Supplementary Figure 1:**
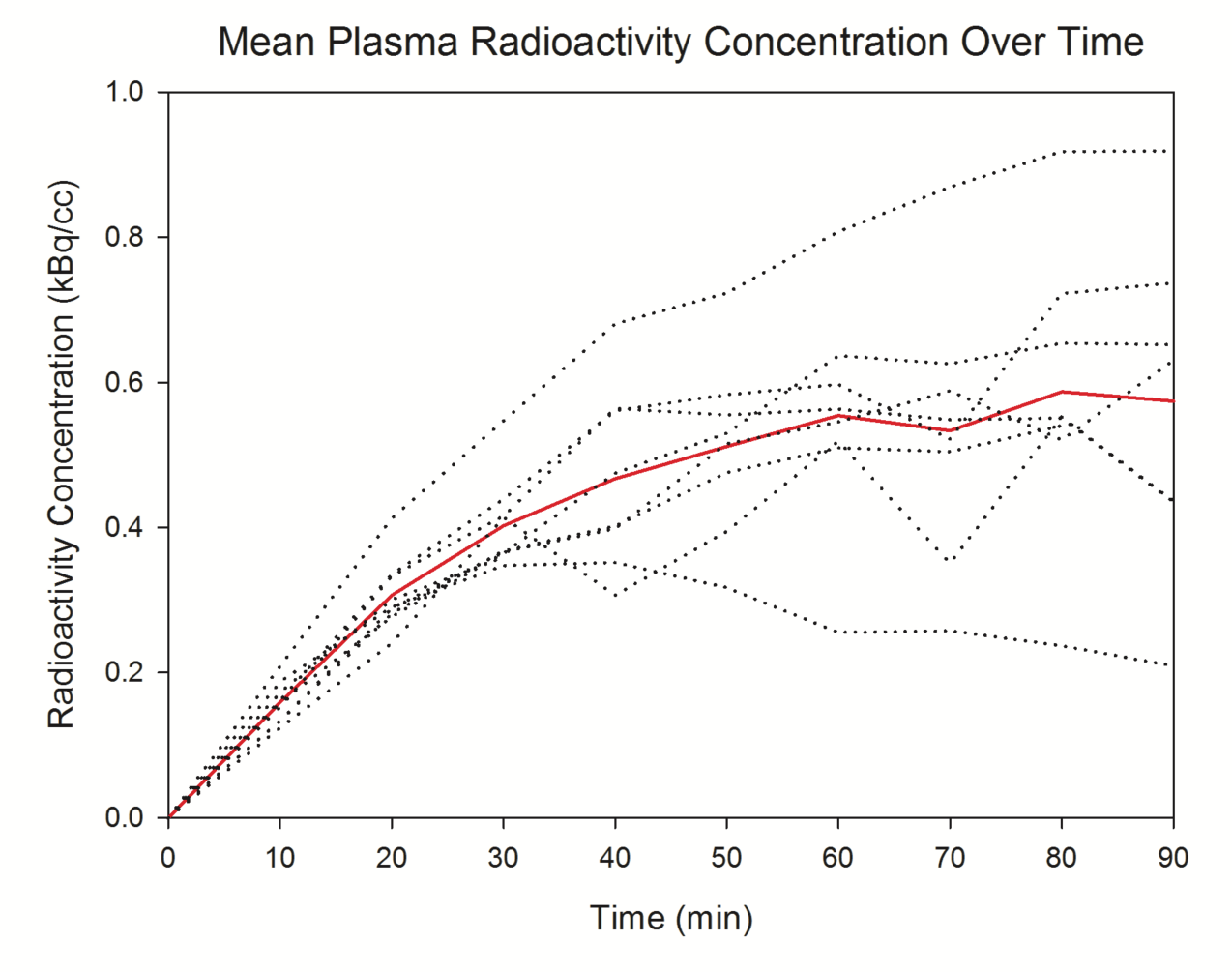
Mean radioactivity concentration (kBq/cc) over the 90min scan. Red line shows the average, dotted lines show individual subject’s concentration over time.

#### ICA Results

In the main text, we reported the results of three separate ICAs: on BOLD-fMRI, static FDG-PET and dynamic FDG-**f**PET; as well as the joint ICA. For each of the 3 separate ICAs, we estimated 7 components and reported the results of the primary visual cortex component in the main text. For the joint analyses, we estimated 3 joint components and reported the results of the first component in the visual cortex. Here, we report the remaining components for each of the ICAs. For clarity, we refer to the components reported in the main text by numerical labels (Component 1, 2, etc.) and components reported in the Supplement by alphabetical labels (Component A, B, etc.).

#### BOLD-fMRI Separate ICA

Supplementary Figure 2 shows the six remaining independent components for the BOLD-fMRI data. The default mode network was obtained in Component A and the dorsal attention network was identified in Component B. The lateral parietal cortex (superior parietal lobule/inferior parietal lobule) superior occipital (superior occipital gyrus, fusiform) and superior temporal cortex was active in Component C. Component D showed activity in higher-visual cortex (cuneus, superior occipital, lingual gyrus) and medial parietal cortex (precuneus) and Component E showed activity in mid cingulate and postcentral gyrus. Component F was dominated by noise, with activity predominantly around the rim of the brain.

In sum, the BOLD-fMRI ICA identified a number of the major networks in the brain, including default mode (component A), dorsal attention (component B), somatosensory (component E) and a number of visual cortex networks (primary visual [Component 1, main text], component C, D). The major noise component identified ‘rim’ effects, which are often associated with subject motion or realignment error (5).

**Supplementary Figure 2:**
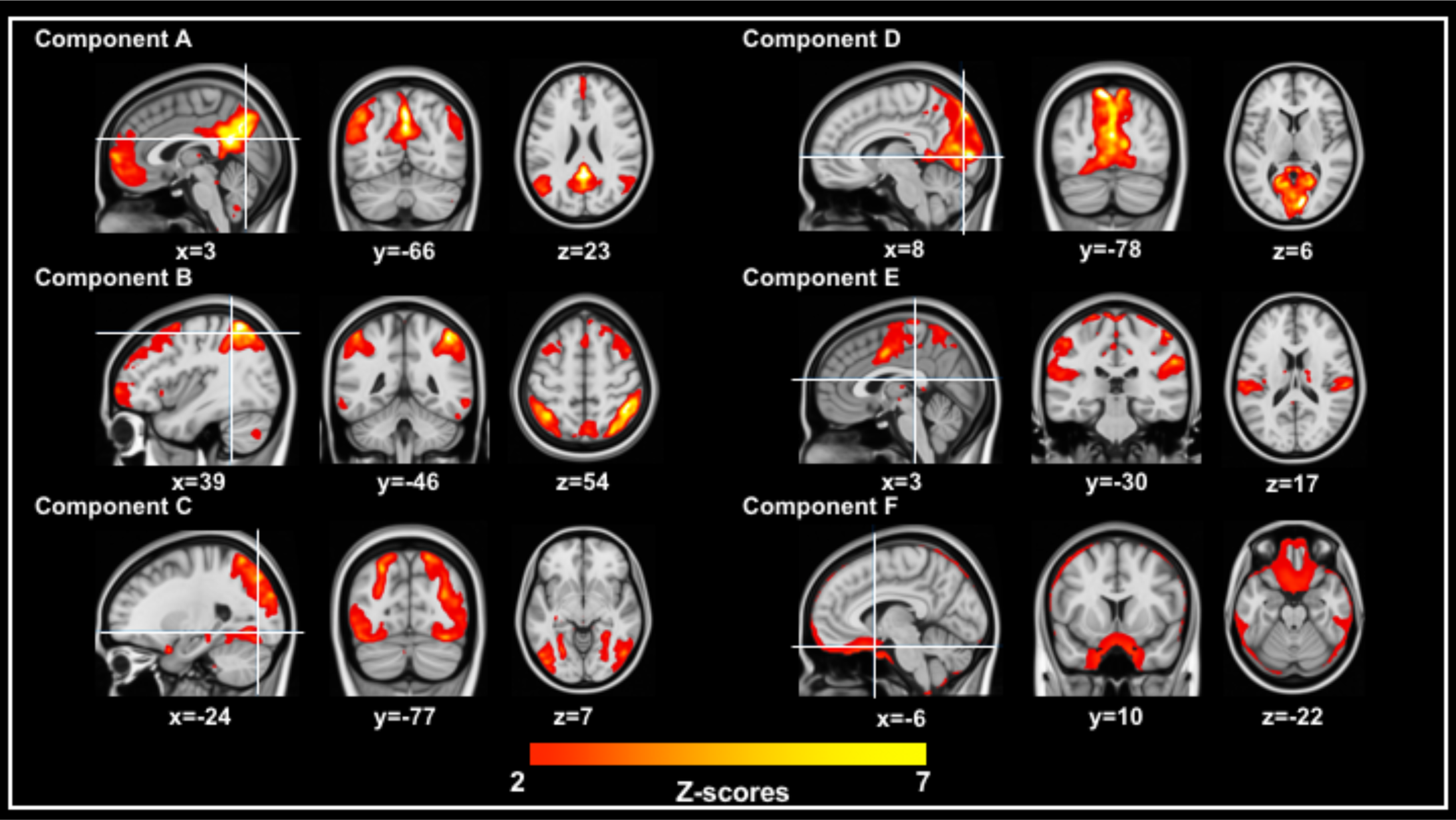
Six BOLD-fMRI components, thresholded to Z=2.0. Coordinates are MNI space.

#### Static FDG-PET Separate ICA

The six static FDG-PET independent components are shown in Supplementary Figure 3. The precuneus and superior medial occipital cortex was predominantly active in Component A, with additional small clusters obtained in frontal (inferior frontal gyrus) and lateral parietal (inferior parietal lobule) regions. Component B showed activity in medial superior frontal and inferior frontal gyrus. Component C showed activity in the cerebellum, and Component D was dominated by noise in the ventricles. Component E showed several small clusters of activity in primarily dorsal regions, encompassing frontal (superior/middle frontal gyrus), central (precentral gyrus, supplementary motor area), parietal (superior parietal lobule) and occipital (superior occipital gyrus) regions. Component F was similar to component D, with more ventral regions active, across frontal (middle/inferior frontal gyrus), central (precentral/postcentral gyrus), parietal (inferior parietal lobule), and occipital (cuneus/middle occipital) and inferior temporal regions.

The static FDG-PET results were qualitatively different from those obtained in BOLD-fMRI. Unlike BOLD-fMRI, were a number of common networks were identified, static FDG-PET showed networks that are not easily reconciled with the BOLD-fMRI literature. The noise component identified here (component D) likely represents the glucose profile of the cerebrospinal fluid (6). It is possible that the cerebellar activity identified in Component C is an artefact of the normalization method used for the sPET analysis. Indeed, we used the mean activity in the cerebellum region to normalize the static PET images for each subject, meaning that the signal in the cerebellum was almost constant across subjects. Components E and F did show a similar activity profile, with component F showing more ventral activity than component D; it may be that these two components represent an underlying metabolic network that has been split across multiple components in this analysis. Further work across larger samples is required to determine the robustness and replicability of these results.

Previous BOLD-fMRI/FDG-PET studies of resting-state connectivity has shown that the metabolic connectome is likely to be characterised by a smaller number of components than BOLD-fMRI (7-9). Di & Biswal (7) and Savio et al. (8) reported reduced anterior-posterior connectivity in resting-state FDG metabolism compared to BOLD-fMRI; with an increase in left-right metabolism between homologous regions in each hemisphere. Those authors conceded that the bolus method may not be sufficient to detect anterior-posterior connectivity, which may occur over a faster timeframe than left-right connectivity. Although we did not specifically test the temporal dynamics of connectivity in this study, our results showing anterior-posterior connectivity in the static FDG analysis (i.e., Components B, E, F) are consistent with this hypothesis. Further work is required to test this hypothesis.

**Supplementary Figure 3:**
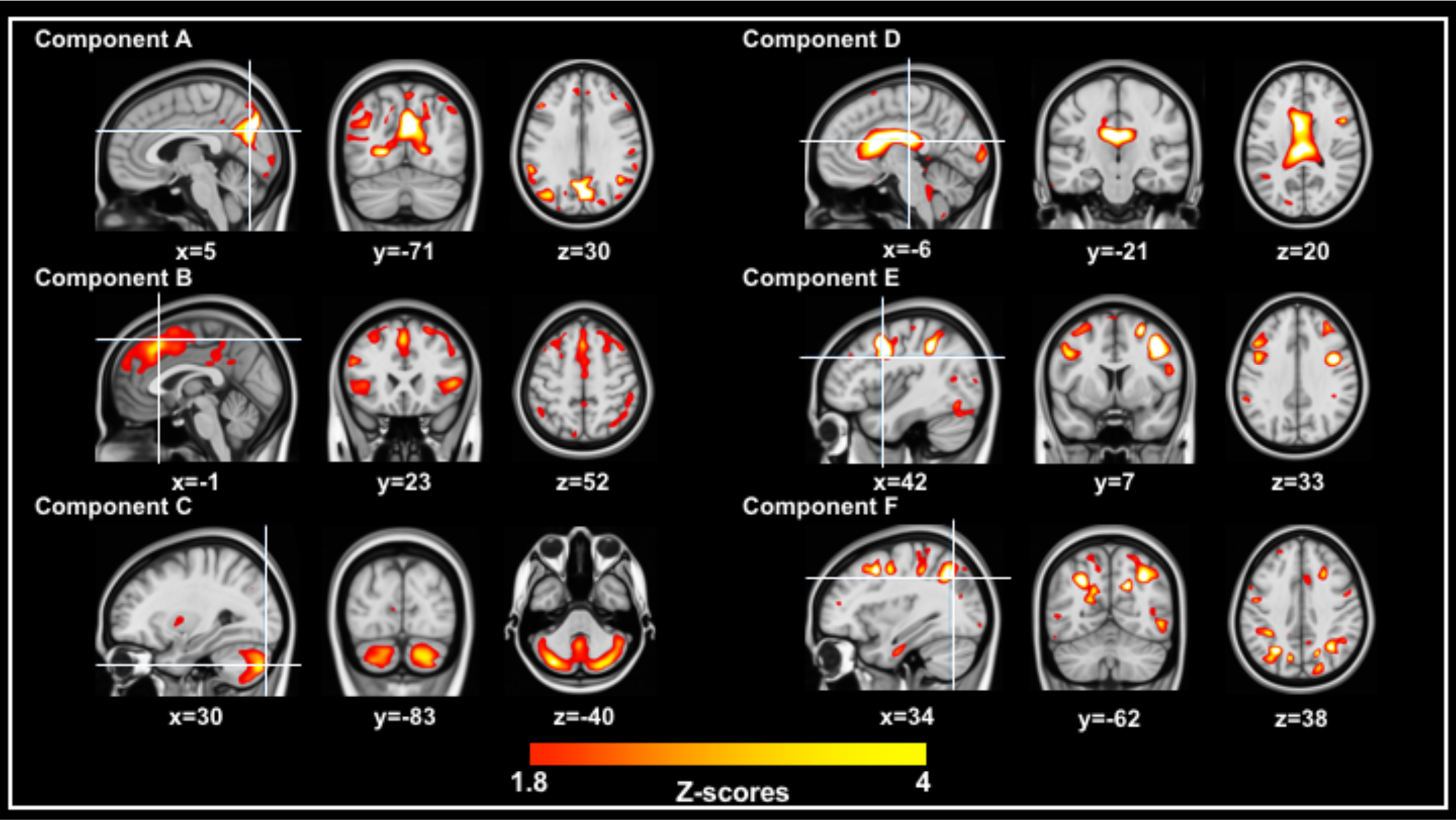
Results of static FDG-PET ICA. Threshold = Z=1.8.

#### Dynamic FDG-fPET Separate ICA

Supplementary Figure 4 shows the results of the separate ICA on FDG-fPET data. As in the previous analyses, ICA estimated 7 components, with the primary component reported in the main text. For the remaining components, Component A was dominated by motor activity in the supplementary motor area and precentral gyrus. Component B showed activity in the superior sagittal, transverse, and straight sinuses. Component D showed small clusters of activity across dorsal cortex. It is unclear if this component represents noise, or is synonymous with the components identified in the static analysis (Supplementary Figure 3; components E & F). Components C, E and F were dominated by noise, showing activity in the ventricles (component C) or disparate clusters across the brain (component E and F).

In sum, the dynamic FDG-fPET analysis identified primary visual (reported in main text) and motor networks. Interestingly, the dynamic analysis identified a venous component (component B) that was not identified in the static analysis. A venous component may indicate global variations in glucose metabolism, identified by fluctuations in the residual FDG draining from the cerebral compartment to the body. It may also be a confound of normalising by the mean grey-matter signal, depicting the temporal dynamics of a tissue to blood FDG ratio.

**Supplementary Figure 4:**
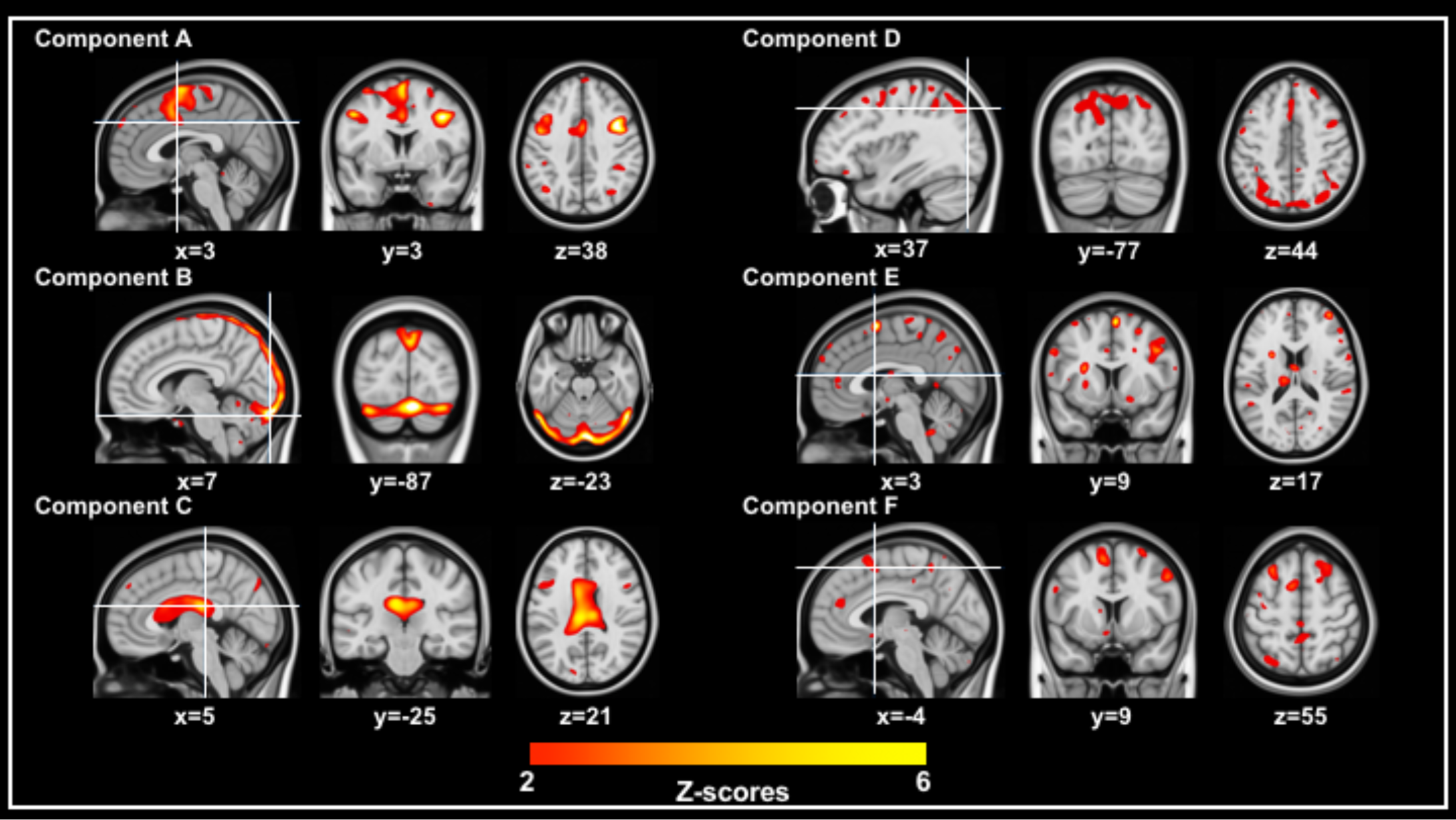
Six FDG-fPET components, thresholded Z=2.0.

#### Joint ICA

We estimated 3 joint components in the ICA; the primary result is reported in text, and the remaining joint components are shown in Supplementary Figure 5. Joint component pair A was dominated by noise – disparate small clusters throughout the brain – in both MR and PET. Joint component pair B showed a relationship between glucose metabolism in the vasculature and BOLD-fMRI activity in the visual cortex.

Joint ICA was conducted on the two primary visual cortex networks identified in the separate ICAs of BOLD-fMRI and FDG-fPET data. This analysis identified a pair of maximally correlated components with activity in the tissue of the primary visual cortex. Interestingly, joint ICA also identified a pair of components showing the glucose metabolism in the sagittal, transverse, and straight sinuses is associated with BOLD activity in the tissue of the visual cortex (Supplementary Figure 5, Component B). This component may be depicting a correlation between downstream BOLD-fMRI effects and reduced venous glucose, both ‘non-local drainage’ responses to neuronal activity. However, caution is advised with this conjecture as the BOLD-fMRI spatial extent is not conclusive, and the FDG-fPET component may be a normalisation artefact as considered in the dynamic FDG-fPET supplementary analysis. Further experiments are required to examine the temporal dynamics of this component to elucidate its origin.

**Supplementary Figure 5:**
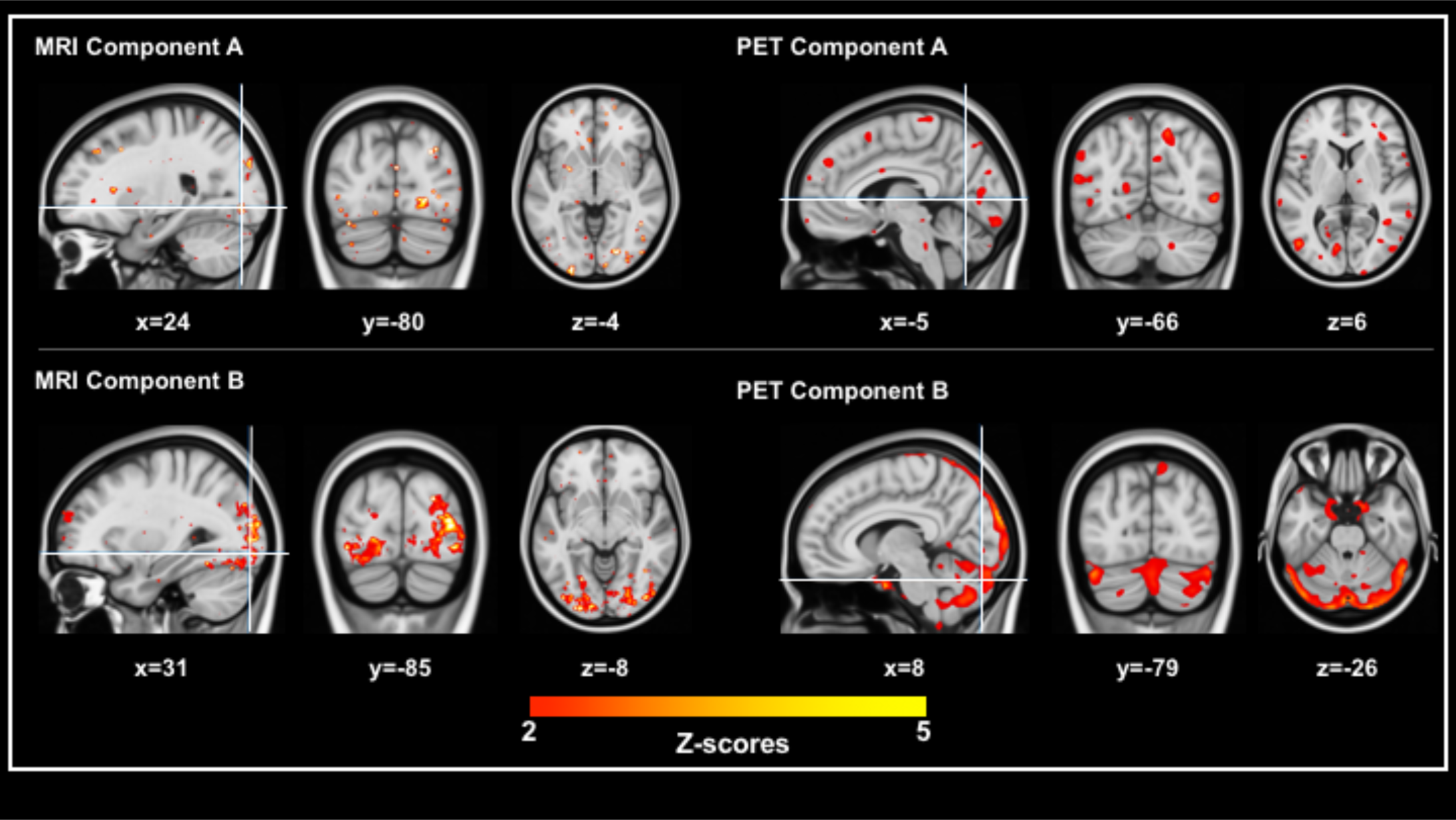
Joint ICA components, threshold Z=2.0.

**Supplementary Table 1:**
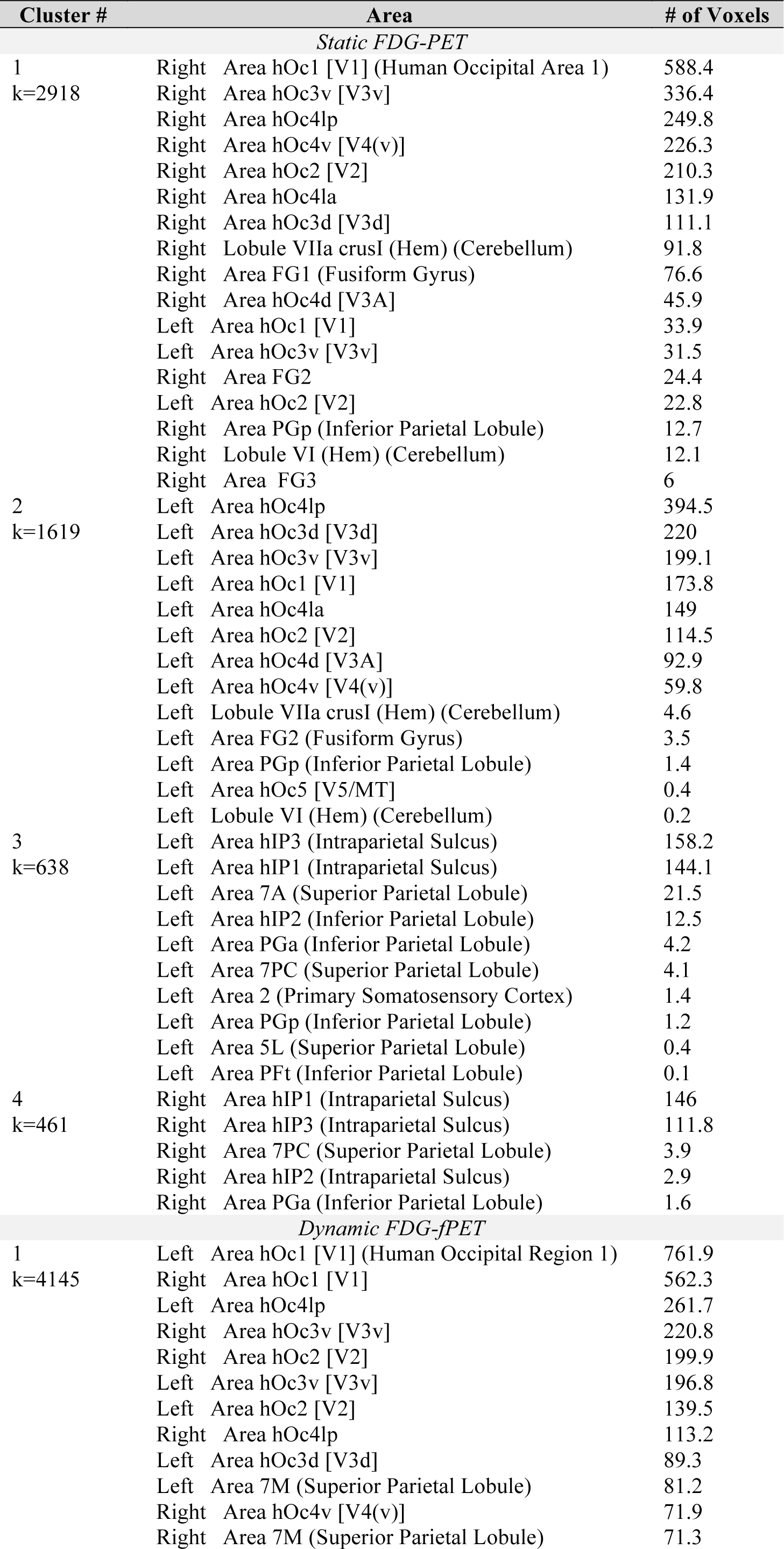
Activated regions in the static FDG-PET, BOLD-fMRI and dynamic FDG-fPET. Component maps for each analysis were thresholded at Z=2, k=200 voxels and entered into a statistical probabilistic anatomical mapping analysis with p<.05. Cluster size (k, number of voxels) is given for each cluster. Note that labels are cytoarchitectonic nomenclature; common region names are given in parentheses where required.

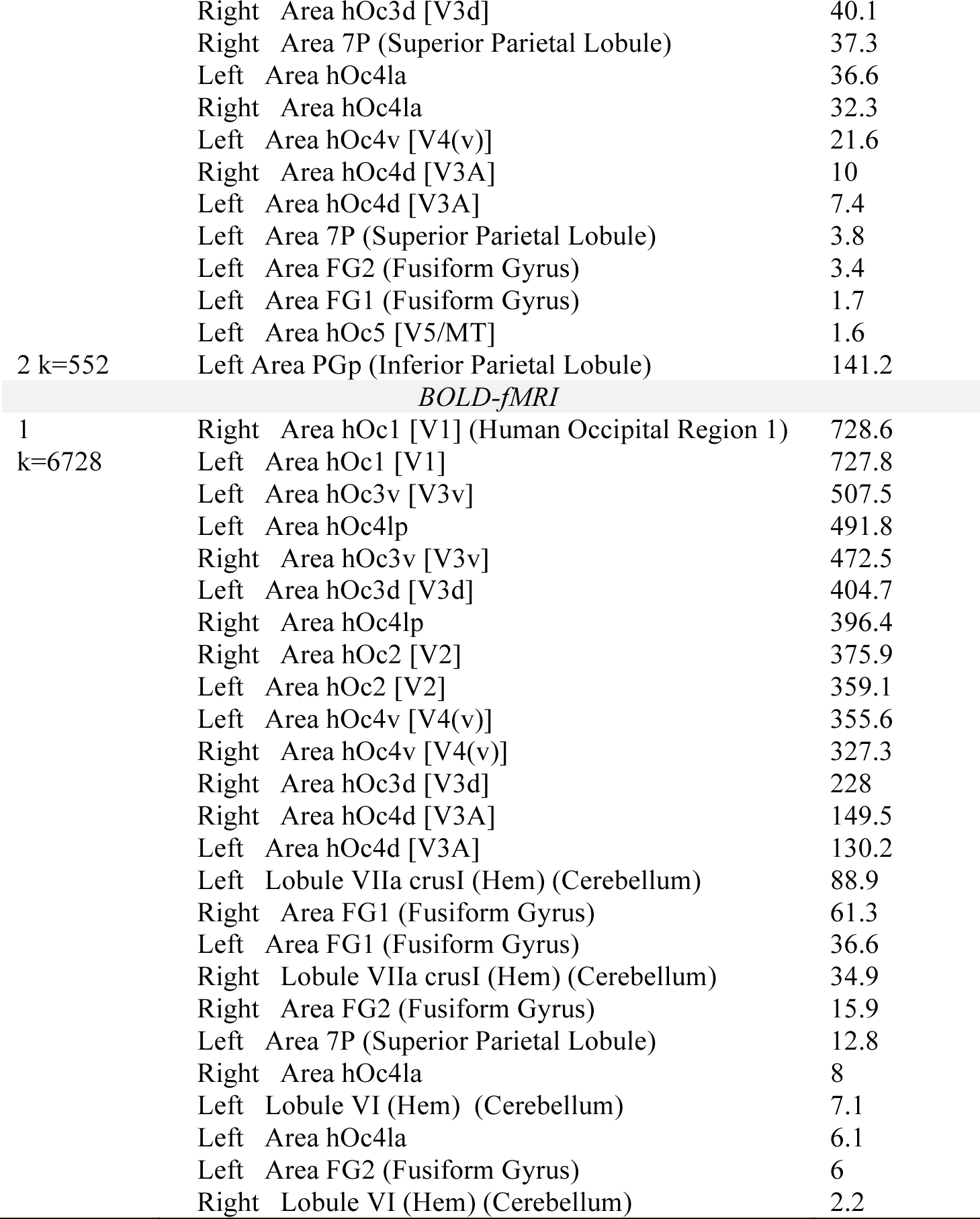

|  |  |  |
| --- | --- | --- |
|  | Right Area hOc3d [V3d] | 40.1 |
|  | Right Area 7P (Superior Parietal Lobule) | 37.3 |
|  | Left Area hOc4la | 36.6 |
|  | Right Area hOc4la | 32.3 |
|  | Left Area hOc4v [V4(v)] | 21.6 |
|  | Right Area hOc4d [V3A] | 10 |
|  | Left Area hOc4d [V3A] | 7.4 |
|  | Left Area 7P (Superior Parietal Lobule) | 3.8 |
|  | Left Area FG2 (Fusiform Gyrus) | 3.4 |
|  | Left Area FG1 (Fusiform Gyrus) | 1.7 |
|  | Left Area hOc5 [V5/MT] | 1.6 |
| 2 k=552 | Left Area PGp (Inferior Parietal Lobule) | 141.2 |
| *BOLD-fMRI* |  |  |
| 1 | Right Area hOc1 [V1] (Human Occipital Region 1) | 728.6 |
| k=6728 | Left Area hOc1 [V1] | 727.8 |
|  | Left Area hOc3v [V3v] | 507.5 |
|  | Left Area hOc4lp | 491.8 |
|  | Right Area hOc3v [V3v] | 472.5 |
|  | Left Area hOc3d [V3d] | 404.7 |
|  | Right Area hOc4lp | 396.4 |
|  | Right Area hOc2 [V2] | 375.9 |
|  | Left Area hOc2 [V2] | 359.1 |
|  | Left Area hOc4v [V4(v)] | 355.6 |
|  | Right Area hOc4v [V4(v)] | 327.3 |
|  | Right Area hOc3d [V3d] | 228 |
|  | Right Area hOc4d [V3A] | 149.5 |
|  | Left Area hOc4d [V3A] | 130.2 |
|  | Left Lobule VIIa crusI (Hem) (Cerebellum) | 88.9 |
|  | Right Area FG1 (Fusiform Gyrus) | 61.3 |
|  | Left Area FG1 (Fusiform Gyrus) | 36.6 |
|  | Right Lobule VIIa crusI (Hem) (Cerebellum) | 34.9 |
|  | Right Area FG2 (Fusiform Gyrus) | 15.9 |
|  | Left Area 7P (Superior Parietal Lobule) | 12.8 |
|  | Right Area hOc4la | 8 |
|  | Left Lobule VI (Hem) (Cerebellum) | 7.1 |
|  | Left Area hOc4la | 6.1 |
|  | Left Area FG2 (Fusiform Gyrus) | 6 |
|  | Right Lobule VI (Hem) (Cerebellum) | 2.2 |

**Supplementary Table 2:** Activated regions in the joint BOLD-fMRI/FDG-fPET analysis. Component maps were thresholded at Z=2, k=200 voxels and entered into a statistical probabilistic anatomical mapping analysis with p<.05. Cluster size (k, number of voxels) is given for each cluster. Note that labels are cytoarchitectonic nomenclature; common region names are given in parentheses where required.

| Cluster # | Area | # of Voxels |
| --- | --- | --- |
| *BOLD-fMRI (Joint Component)* |  |  |
| 1<br>k=6116 | Right Area hOc1 [V1] (Human Occipital Region 1) | 605.7 |
|  | Left Area hOc4lp | 591.6 |
|  | Left Area hOc1 [V1] | 582.6 |
|  | Left Area hOc3v [V3v] | 484.6 |
|  | Right Area hOc4lp | 389.3 |
|  | Right Area hOc3v [V3v] | 366.2 |
|  | Right Area hOc2 [V2] | 328.8 |
|  | Left Area hOc2 [V2] | 317.6 |
|  | Left Area hOc3d [V3d] | 304.6 |
|  | Left Area hOc4v [V4(v)] | 265.4 |
|  | Right Area hOc3d [V3d] | 206.6 |
|  | Right Area hOc4v [V4(v)] | 192 |
|  | Right Area hOc4d [V3A] | 138.4 |
|  | Left Area hOc4d [V3A] | 102.3 |
|  | Right Area hOc4la | 88.6 |
|  | Left Area hOc4la | 80.1 |
|  | Left Lobule VIIa crusI (Hem) (Cerebellum) | 67.3 |
|  | Right Area FG1 (Fusiform Gyrus) | 65.2 |
|  | Left Lobule VI (Hem) (Cerebellum) | 30 |
|  | Right Area FG2 (Fusiform Gyrus) | 30 |
|  | Left Area FG1 (Fusiform Gyrus) | 21.6 |
|  | Right Area FG4 (Fusiform Gyrus) | 19.2 |
|  | Right Lobule VI (Hem) (Cerebellum) | 9.5 |
|  | Right Lobule VIIa crusI (Hem) (Cerebellum) | 8.8 |
|  | Left Area FG2 (Fusiform Gyrus) | 8.2 |
| *FDG-fPET (Joint Component)* |  |  |
| 1<br>k=3032 | Right Area hOc1 [V1] (Human Occipital Region 1) | 639.7 |
|  | Left Area hOc1 [V1] | 480.6 |
|  | Right Area hOc3v [V3v] | 249.5 |
|  | Right Area hOc2 [V2] | 241.5 |
|  | Left Area hOc3v [V3v] | 184.8 |
|  | Right Area hOc4lp | 176.2 |
|  | Left Area hOc4lp | 166.2 |
|  | Left Area hOc2 [V2] | 90.3 |
|  | Left Area hOc4d [V3A] | 59.9 |
|  | Right Area hOc4d [V3A] | 59.5 |
|  | Left Area hOc4la | 55.6 |
|  | Left Area hOc3d [V3d] | 53.2 |
|  | Right Area hOc3d [V3d] | 26.2 |
|  | Left Area hOc4v [V4(v)] | 22.9 |
|  | Right Area hOc4v [V4(v)] | 18 |
|  | Left Lobule VI (Hem) (Cerebellum) | 14.4 |
|  | Right Area hOc4la | 13.8 |
|  | Left Area FG1 (Fusiform Gyrus) | 0.7 |
|  | Left Lobule VI (Verm) (Cerebellum) | 0.5 |

## Abbreviations

BOLD-fMRI/FDG-PET: simultaneously acquired blood oxygenation level dependent functional magnetic resonance imaging/ fluorodeoxyglucose positron emission tomography
BSL: blood sugar level
SPAM: statistical parametric anatomical mapping

## References

1. Mergenthaler P, Lindauer U, Dienel GA, & Meisel A (2013) Sugar for the brain: the role of glucose in physiological and pathological brain function. Trends in neurosciences 36(10): 587–597.

2 Serlin Y, Shelef I, Knyazer B, & Friedman A (2015) Anatomy and physiology of the blood–brain barrier. Seminars in cell & developmental biology, (Elsevier), pp 2–6.

3 Kety SS (1957) The general metabolism of the brain in vivo. Metabolism of the nervous system, (Elsevier), pp 221–237.

4. Sokoloff L (1960) The metabolism of the central nervous system in vivo. Handbook of Physiology, Section I, Neurophysiology 3: 1843–1864.

5. Harris JJ, Jolivet R, & Attwell D (2012) Synaptic energy use and supply. Neuron 75(5): 762–777.

6. Petit-Taboue M, Landeau B, Desson J, Desgranges B, & Baron J (1998) Effects of healthy aging on the regional cerebral metabolic rate of glucose assessed with statistical parametric mapping. Neuroimage 7(3): 176–184.

7. Mosconi L, et al. (2009) FDG-PET changes in brain glucose metabolism from normal cognition to pathologically verified Alzheimer’s disease. European journal of nuclear medicine and molecular imaging 36(5): 811–822.

8. Pagano G, Niccolini F, & Politis M (2016) Current status of PET imaging in Huntington’s disease. European journal of nuclear medicine and molecular imaging 43(6): 1171–1182.

9 Moses WW (2011) Fundamental limits of spatial resolution in PET. Nuclear Instruments and Methods in Physics Research Section A: Accelerators, Spectrometers, Detectors and Associated Equipment 648:S236-S240.

10. Feinberg DA & Yacoub E (2012) The rapid development of high speed, resolution and precision in fMRI. Neuroimage 62(2): 720–725.

11. Yacoub E, et al. (2001) Imaging brain function in humans at 7 Tesla. Magnetic Resonance in Medicine 45(4): 588–594.

12. Logothetis NK, Pauls J, Augath M, Trinath T, & Oeltermann A (2001) Neurophysiological investigation of the basis of the fMRI signal. Nature 412(6843): 150.

13. Logothetis NK (2008) What we can do and what we cannot do with fMRI. Nature 453(7197): 869.

14 Tong Y, Yao J, Chen JJ, & Frederick Bd (2018) The resting-state fMRI arterial signal predicts differential blood transit time through the brain. Journal of Cerebral Blood Flow & Metabolism:0271678X17753329.

15 Chen Z, et al. (2018) From simultaneous to synergistic MR- PET brain imaging: A review of hybrid MR- PET imaging methodologies. Human brain mapping.

16. Wehrl HF, et al. (2013) Simultaneous PET-MRI reveals brain function in activated and resting state on metabolic, hemodynamic and multiple temporal scales. Nature medicine 19(9): 1184.

17. Riedl V, et al. (2014) Local activity determines functional connectivity in the resting human brain: a simultaneous FDG-PET/fMRI study. Journal of neuroscience 34(18): 6260–6266.

18. Carson RE (2000) PET physiological measurements using constant infusion. Nuclear medicine and biology 27(7): 657–660.

19. Villien M, et al. (2014) Dynamic functional imaging of brain glucose utilization using fPET-FDG. Neuroimage 100: 192–199.

20. Hahn A, et al. (2016) Quantification of task-specific glucose metabolism with constant infusion of 18F-FDG. Journal of Nuclear Medicine 57(12): 1933–1940.

21. Hahn A, et al. (2018) Task-relevant brain networks identified with simultaneous PET/MR imaging of metabolism and connectivity. Brain Structure and Function 223(3): 1369–1378.

22. Rischka L, et al. (2018) Reduced task durations in functional PET imaging with [18F] FDG approaching that of functional MRI. NeuroImage 181: 323–330.

23. Burgos N, et al. (2014) Attenuation correction synthesis for hybrid PET-MR scanners: application to brain studies. IEEE transactions on medical imaging 33(12): 2332–2341.

24. Sforazzini F, et al. (2017) MR-based attenuation map re-alignment and motion correction in simultaneous brain MR-PET imaging. Biomedical Imaging (ISBI 2017), 2017 IEEE 14th International Symposium on, (IEEE), pp 231–234.

25. Panin VY, Kehren F, Michel C, & Casey M (2006) Fully 3-D PET reconstruction with system matrix derived from point source measurements. IEEE transactions on medical imaging 25(7): 907–921.

26 Beckmann CF, DeLuca M, Devlin JT, & Smith SM (2005) Investigations into resting-state connectivity using independent component analysis. Philosophical Transactions of the Royal Society B: Biological Sciences 360(1457):1001–1013.

27. Calhoun VD, Adali T, Pearlson GD, & Pekar J (2001) A method for making group inferences from functional MRI data using independent component analysis. Human brain mapping 14(3): 140–151.

28. Avants BB, et al. (2011) A reproducible evaluation of ANTs similarity metric performance in brain image registration. Neuroimage 54(3): 2033–2044.

29. Avants BB, et al. (2010) The optimal template effect in hippocampus studies of diseased populations. Neuroimage 49(3): 2457–2466.

30 Bauer CM, Cabral H, Greve D, & Killiany R (2013) Differentiating between normal aging, mild cognitive impairment, and Alzheimer’s disease with FDG-PET: Effects of normalization region and partial volume correction method. J Alzheimers Dis Parkinsonism 3(1).

31 Hyvärinen A & Oja E (2000) Independent component analysis: algorithms and applications. Neural networks 13(4–5):411–430.

32 Beckmann CF, Mackay CE, Filippini N, & Smith SM (2009) Group comparison of resting-state FMRI data using multi-subject ICA and dual regression. Neuroimage 47(Suppl 1):S148.

33. Filippini N, et al. (2009) Distinct patterns of brain activity in young carriers of the APOE-∊4 allele. Proceedings of the National Academy of Sciences 106(17): 7209–7214.

34. Calhoun VD, Adali T, Pearlson G, & Kiehl K (2006) Neuronal chronometry of target detection: fusion of hemodynamic and event-related potential data. Neuroimage 30(2): 544–553.

35. Lee T-W, Girolami M, & Sejnowski TJ (1999) Independent component analysis using an extended infomax algorithm for mixed subgaussian and supergaussian sources. Neural computation 11(2): 417–441.

36. Eickhoff SB, et al. (2005) A new SPM toolbox for combining probabilistic cytoarchitectonic maps and functional imaging data. Neuroimage 25(4): 1325–1335.

37. Eickhoff SB, Heim S, Zilles K, & Amunts K (2006) Testing anatomically specified hypotheses in functional imaging using cytoarchitectonic maps. Neuroimage 32(2): 570–582.

38. Ward PG, et al. (2018) Combining images and anatomical knowledge to improve automated vein segmentation in MRI. NeuroImage 165: 294–305.

39. Di X, Biswal, & Alzheimer’s Disease Neuroimaging Initiative BB (2012) Metabolic brain covariant networks as revealed by FDG-PET with reference to resting-state fMRI networks. Brain connectivity 2(5): 275–283.

40. Savio A, et al. (2017) Resting-State Networks as Simultaneously Measured with Functional MRI and PET. Journal of Nuclear Medicine 58(8): 1314–1317.

41. Calhoun VD, Kiehl KA, & Pearlson GD (2008) Modulation of temporally coherent brain networks estimated using ICA at rest and during cognitive tasks. Human brain mapping 29(7): 828–838.

42. Calhoun V, et al. (2006) Method for multimodal analysis of independent source differences in schizophrenia: combining gray matter structural and auditory oddball functional data. Human brain mapping 27(1): 47–62.

43. Calhoun VD & Adali T (2009) Feature-based fusion of medical imaging data. IEEE Transactions on Information Technology in Biomedicine 13(5): 711–720.

44. Tomasi DG, et al. (2017) Dynamic brain glucose metabolism identifies anti-correlated cortical-cerebellar networks at rest. Journal of Cerebral Blood Flow & Metabolism 37(12): 3659–3670.

45. Aiello A, Banzer P, Neugebauer M, & Leuchs G (2015) From transverse angular momentum to photonic wheels. Nature Photonics 9(12): 789.

46. Tomasi D, Wang G-J, & Volkow ND (2013) Energetic cost of brain functional connectivity. Proceedings of the National Academy of Sciences 110(33): 13642–13647.

47. Catana C, et al. (2011) MRI-assisted PET motion correction for neurologic studies in an integrated MR-PET scanner. Journal of Nuclear Medicine 52(1): 154–161.

48. Shah N, et al. (2017) Multimodal fingerprints of resting state networks as assessed by simultaneous trimodal MR-PET-EEG imaging. Scientific reports 7(1): 6452.

49. Gjedde A, Kuwabara H, & Hakim AM (1990) Reduction of functional capillary density in human brain after stroke. Journal of Cerebral Blood Flow & Metabolism 10(3): 317–326.

50. Gould IG, Tsai P, Kleinfeld D, & Linninger A (2017) The capillary bed offers the largest hemodynamic resistance to the cortical blood supply. Journal of Cerebral Blood Flow & Metabolism 37(1): 52–68.

51 Bright MG, Croal PL, Blockley NP, & Bulte DP (2017) Multiparametric measurement of cerebral physiology using calibrated fMRI. NeuroImage.

52. Calhoun VD & Sui J (2016) Multimodal fusion of brain imaging data: A key to finding the missing link (s) in complex mental illness. Biological psychiatry: cognitive neuroscience and neuroimaging 1(3): 230–244.

53. Jamadar S, Hughes M, Fulham WR, Michie PT, & Karayanidis F (2010) The spatial and temporal dynamics of anticipatory preparation and response inhibition in task-switching. Neuroimage 51(1): 432–449.

54 Group F-NBW (2016) BEST (biomarkers, endpoints, and other tools) resource.

55. Biswal B, Zerrin Yetkin F, Haughton VM, & Hyde JS (1995) Functional connectivity in the motor cortex of resting human brain using echo- planar mri. Magnetic resonance in medicine 34(4): 537–541.

## References

1. Hahn A, et al. (2016) Quantification of task-specific glucose metabolism with constant infusion of 18F-FDG. J Nucl Med 57(12): 1933–1940.

2. Villien M, et al. (2014) Dynamic functional imaging of brain glucose utilization using fPET-FDG. Neuroimage 100: 192–199.

3 Bailey DL, Townsend DW, Valk PE, & Maisey MN (2005) Positron emission tomography (Springer).

4. Carson RE, et al. (1993) Comparison of bolus and infusion methods for receptor quantitation: application to [18F] cyclofoxy and positron emission tomography. J Cereb Blood Flow Metab 13(1): 24–42.

5. Diedrichsen J & Shadmehr R (2005) Detecting and adjusting for artifacts in fMRI time series data. Neuroimage 27(3): 624–634.

6. Spector R, Snodgrass SR, & Johanson CE (2015) A balanced view of the cerebrospinal fluid composition and functions: focus on adult humans. Exp Neurol 273: 57–68.

7. Di X, Biswal, & Alzheimer’s Disease Neuroimaging Initiative BB (2012) Metabolic brain covariant networks as revealed by FDG-PET with reference to resting-state fMRI networks. Brain Connect 2(5): 275–283.

8. Savio A, et al. (2017) Resting-State Networks as Simultaneously Measured with Functional MRI and PET. J Nucl Med 58(8): 1314–1317.

9. Tomasi DG, et al. (2017) Dynamic brain glucose metabolism identifies anti-correlated cortical-cerebellar networks at rest. J Cereb Blood Flow Metab 37(12): 3659–3670.

